# Super-resolution single molecule network analysis (SuperResNET) detects changes to clathrin structure by small molecule inhibitors

**DOI:** 10.1101/2024.03.07.583946

**Authors:** Timothy H. Wong, Ismail M. Khater, Christian Hallgrimson, Y. Lydia Li, Ghassan Hamarneh, Ivan R. Nabi

**Affiliations:** Department of Cellular & Physiological Sciences, Life Sciences Institute, University of British Columbia, Vancouver, BC, Canada V6T 1Z3; School of Computing Science, Simon Fraser University, Burnaby, BC, Canada V5A 1S6; Department of Electrical and Computer Engineering, Faculty of Engineering and Technology, Birzeit University, Birzeit P627, Palestine; School of Biomedical Engineering, University of British Columbia, Vancouver, BC, Canada V6T 1Z3

## Abstract

Specificity of small molecules for their target molecule in the cell is critical to determine their effective use as biologics and therapeutics. Small molecule inhibitors of clathrin endocytosis, Pitstop 2, and the dynamin inhibitor Dynasore, have off-target effects and their specificity has been challenged. Here, we used SuperResNET to apply network analysis to 20 nm resolution dSTORM single-molecule localization microscopy (SMLM) to test whether Pitstop 2 and Dynasore alter the morphology of clathrin coated pits in intact cells. SuperResNET analysis of dSTORM data from HeLa and Cos7 cells identifies three classes of clathrin structures: small oligomers (Class I); pits and vesicles (Class II); and larger clusters corresponding to fused clathrin pits and clathrin plaques (Class III). SuperResNET analysis of high resolution MinFlux imaging identifies Class 1 oligomers as well as Class 2 structures including morphologically identifiable clathrin pits and vesicles. SuperResNET feature analysis of dSTORM data shows that Pitstop 2 and Dynasore induce the formation of distinct homogenous populations of clathrin structures in HeLa cells. Pitstop 2 blobs are smaller and more elongated than those induced by Dynasore, indicating that these two clathrin inhibitors arrest clathrin endocytosis at distinct stages. Pitstop 2 and Dynasore are not impacting clathrin structure via actin depolymerization as the actin depolymerizing agent latrunculin A (LatA) induced larger heterogeneous clathrin structures. Ternary analysis of SuperResNET shape features presents a distinct profile for Pitstop 2 Class II structures. The most representative Pitstop blobs align with and resemble MinFlux clathrin pits while control structures resemble Minflux clathrin vesicles. SuperResNET analysis of SMLM data is therefore a highly sensitive approach to detect the effect of small molecules on target molecule structure in situ in the cell.

## Introduction

Small molecules generated by medicinal chemistry (i.e. man-made) as well as from natural products are important tools in both basic biology discoveries and as clinical therapeutics [1, 2]. Large compound small molecule libraries are often screened for biological activity and efficacy by high-throughput screening for specific effects on target molecules and cellular functions. Critical to the use of these drugs are target specificity, biological activity and efficacy [3]. Key to validation of specificity is whether the small molecule targets the molecule of interest. Indeed, many small molecules are insufficiently characterized and are later found to be nonselective [4]. Different approaches using high throughput and functional screens have been developed to identify and characterize target specificity of these drugs [5]. CRISPR screens can determine how genetic deletions, knockdowns and overexpression effect drug sensitivity [6]. However, these approaches do not report on whether the small molecule actually targets and impacts the target molecule within the cell. Structural biology approaches, such as X-ray crystallography and cryoEM of the drug target complex, can demonstrate direct interaction between the drug and target protein [7]. While structural biology approaches provide angstrom resolution, these interactions occur ex vivo and do not report on small molecule-target interactions in the cell.

Whole cell super-resolution microscopy approaches with selective labeling is a powerful approach for structural analysis of molecular and macromolecular structures in the mesoscale range of 10-100nm [8]. Single molecule localization microscopy (SMLM) approaches such as direct stochastic optical reconstruction microscopy (dSTORM), localizes single fluorophores with 20-30 nm XY and 50-60 nm Z resolution [9–11]. A more recent SMLM approach, MinFlux, applies a donut shaped excitation laser, as for simulated emission depletion (STED) microscopy, to SMLM to reach 2 nm isotropic resolution [12]. As opposed to pixel-based images generated by conventional microscopy, SMLM produces an event list of localizations, also known as point cloud, highly suited to the application of computational imaging, network modeling, and machine learning based 3D pattern analysis tools. The combination of super-resolution microscopy with computational and machine learning tools have been proposed to be promising tools for advancing drug development and nanomedicines [13, 14].

SuperResNET is a network analysis pipeline for the analysis of point cloud data generated by SMLM. The analysis pipeline processes the point cloud data to generate quantitative data of protein clusters and visualizes representative clusters. SuperResNET is equipped with computational modules for iterative merging, filtering, segmenting clusters of localizations, extracting various cluster features, and applying classification to identify the biological clusters [15]. Previously, we applied SuperResNET to caveolin-1 (Cav1) point clouds to classify caveolae and smaller non-caveolar oligomeric structures called scaffolds [15, 16]; importantly, we were able to show that SuperResNET analysis of 3D dSTORM Cav1 datasets could detect structural changes induced by F92A/V94A point mutations to the Cav1 scaffolding domain [17]. This highlights the ability of SMLM network analysis to detect structural changes of macromolecular structures in situ within the cell.

To test whether SuperResNET could detect molecular changes to target proteins induced by small molecule inhibitors, we studied the well-characterized clathrin endocytic pathway. Clathrin light chain and heavy chain coat proteins are recruited to the site of clathrin endocytosis, where they interact with adaptor proteins and recruit cargo to nascent clathrin oligomers; the clathrin triskelion induces membrane invagination to form coated pits that upon dynamin and bar domain proteins mediated membrane scission release clathrin coated vesicles from the plasma membrane [18]. Pitstop 2, a small molecule inhibitor of clathrin endocytosis, binds to the N-terminal domain of clathrin heavy chain, preventing interaction with clathrin-box motif-containing peptides and blocking clathrin endocytosis [19]. However, no significant changes in clathrin pit structure were observed by EM and specificity of Pitstop 2 for clathrin has been challenged as it also inhibits clathrin-independent endocytosis of CD44, CD98 and CD147 [19, 20]. Dynasore, a non-competitive inhibitor of dynamin 1 and dynamin 2, rapidly blocks clathrin coated vesicle formation [21]. However, Dynasore addition to triple dynamin KO fibroblast also disrupts F-actin, showing that it has off-target effects unrelated to dynamin [22].

Here, using SuperResNET SMLM network analysis, we report that Pitstop 2 and Dynasore both induce the formation of homogenous populations of clathrin structures distinct from the larger heterogeneous structures induced by the actin depolymerizing agent latrunculin A (LatA). This indicates that the two clathrin inhibitors arrest clathrin pit maturation and that their activity is not mediated by actin depolymerization. Clathrin structures induced by Pitstop 2 were smaller and more elongated than those induced by Dynasore suggesting that these two clathrin endocytosis inhibitors arrest endocytosis at different stages of clathrin coat formation. Shape feature comparison with clathrin structures analyzed by MinFlux indicates that Pitstop arrests clathrin pit maturation. Detection of changes to target clathrin structures by small molecule inhibitors of clathrin endocytosis shows that SuperResNET analysis represents a highly sensitive approach to detect changes in molecular structure to targets of small molecule inhibitors in situ in the cell.

## Results

### SuperResNET SMLM network analysis detects three classes of clathrin structures

HeLa cells were labeled with anti-clathrin heavy chain primary and Alexa647 conjugated Fab_2_ secondary antibodies and imaged by 3D TIRF dSTORM. The presence of multiple hollow clathrin coated pits can be observed in a pixel rendered image of the dSTORM data (Fig. 1A). The 3D point cloud event list generated by dSTORM was processed using SuperResNET, SMLM network analysis software that performs iterative merging, filtering, segmentation and classification [15]. Cluster features, 29 description vectors encompassing 3D size, volume/area, topology and hollowness, shape and node network features for the clusters/blobs. The extracted features are then inputted to the K-means clustering method to discern 3 distinct classes. The cluster means derived from these features can subsequently be utilized to categorize additional blobs from diverse cells into one of the three identified classes. Fig 1B shows a representative image of the same cell rendered after being processed by SuperResNET. A selected ROI of that image shows segmentation and classification of individual clathrin structures into three groups, smaller (colored red), intermediate (green) and larger (blue) clusters (Fig. 1C). Z-stack representation of a number of coated pits from the ROI shows that hollow pits extend over 100-150 nm in Z and correspond with blobs that are classified as Class II structures (Fig. 1D). The batch analysis pipeline of SuperResNET consists of computational modules to process SMLM event lists and output quantifiable class features and visualization of blobs to analyze different control and treatment groups across different replicates and experiments (Fig. 1E).

**Figure 1.**
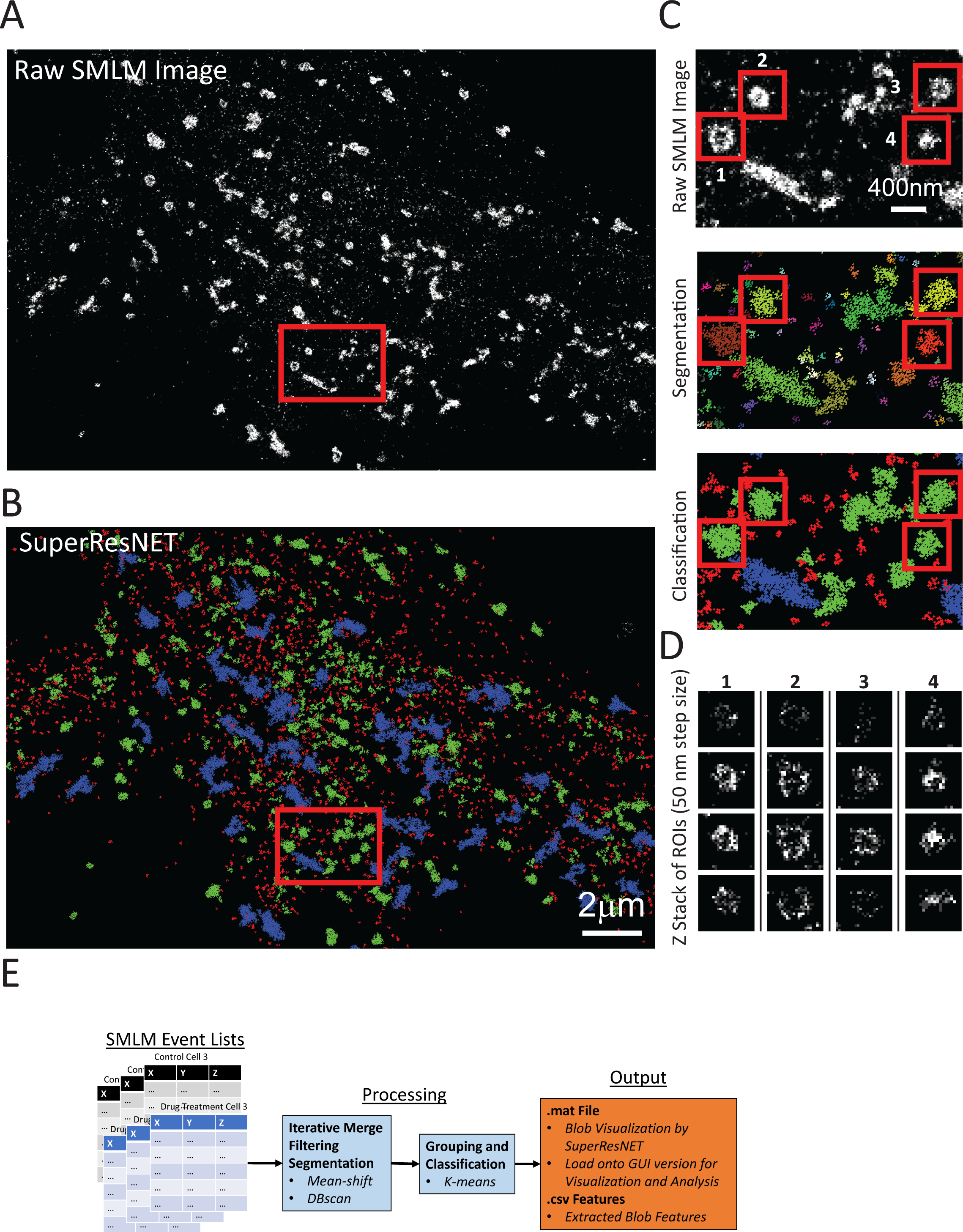
SuperResNET segmentation and classification of clathrin clusters from 3D SMLM data. (A) Representative 3D SMLM dSTORM image of clathrin heavy chain antibody labeling in a HeLa cell. Scale bar 2 µm. (B) Corresponding image processed by SuperResNET showing clathrin clusters segmented as Class I (red), Class II (green) and Class III (blue). Scale bar 2 µm. (C) ROI showing a pixel representation of the raw event list (20 nm pixel size per localization), SuperResNet DBSCAN cluster segmentation and grouping into three classes based on feature analysis. (D) 50 nm thickness Z stack of hollow clathrin pits highlighted in C (red squares). (E) SuperResNET batch analysis pipeline consisting of modules to process SMLM event lists from multiple experiments with different treatment groups. Pipeline outputs .CSV that can be used to quantify blob features and .MAT cell and segmented blob files that can be used for blob visualization or other analyzes with SuperResNET GUI version.

To assess robustness of SuperResNET group classification of clathrin in HeLa cells, we applied the same group classification to clathrin labeled Cos7 cells (Fig. 2A). Quantification of the proportion of each class showed similar distributions of the three classes between HeLa and Cos7 cells, with the vast majority of clusters consisting of the smaller red class and a minority consisting of the larger blue class (Fig 2B). Feature analysis of the three classes in both cell lines identify Class I as small clusters of ∼75 nm diameter (average X range), Class II as intermediate clusters with a diameter of ∼300 nm and Class III as larger structures of ∼800 nm diameter. Class II blobs are more spherical than Class I or III blobs; considering primary and secondary antibody length and dSTORM microscopy resolution of ∼20 nm, the diameter of Class II blobs correspond most closely with 100-200 nm diameter clathrin pits and vesicles (Fig 2C) [23, 24]. Consistently, most hollow blobs identified by SMLM in Figure 1 are classified as Class II blobs. Class III are much larger in both X and Z range (Fig 2C) and include blobs that are not effectively segmented from adjacent structures; this class may also include larger clathrin plaques (Supp. Fig. 1) [25, 26]. Euclidean distance analysis for 27 features (excluding volume and area), showed that Class I and Class II are highly similar between HeLa and Cos7 cells (Supp. Fig. 2), highlighting similarities between these two classes of clathrin structures between the two cell types. The large variability of Class III blobs between the two cell lines may reflect challenges in segmenting closely opposed clusters, suggesting that these blobs do not represent distinct clathrin states.

**Figure 2.**
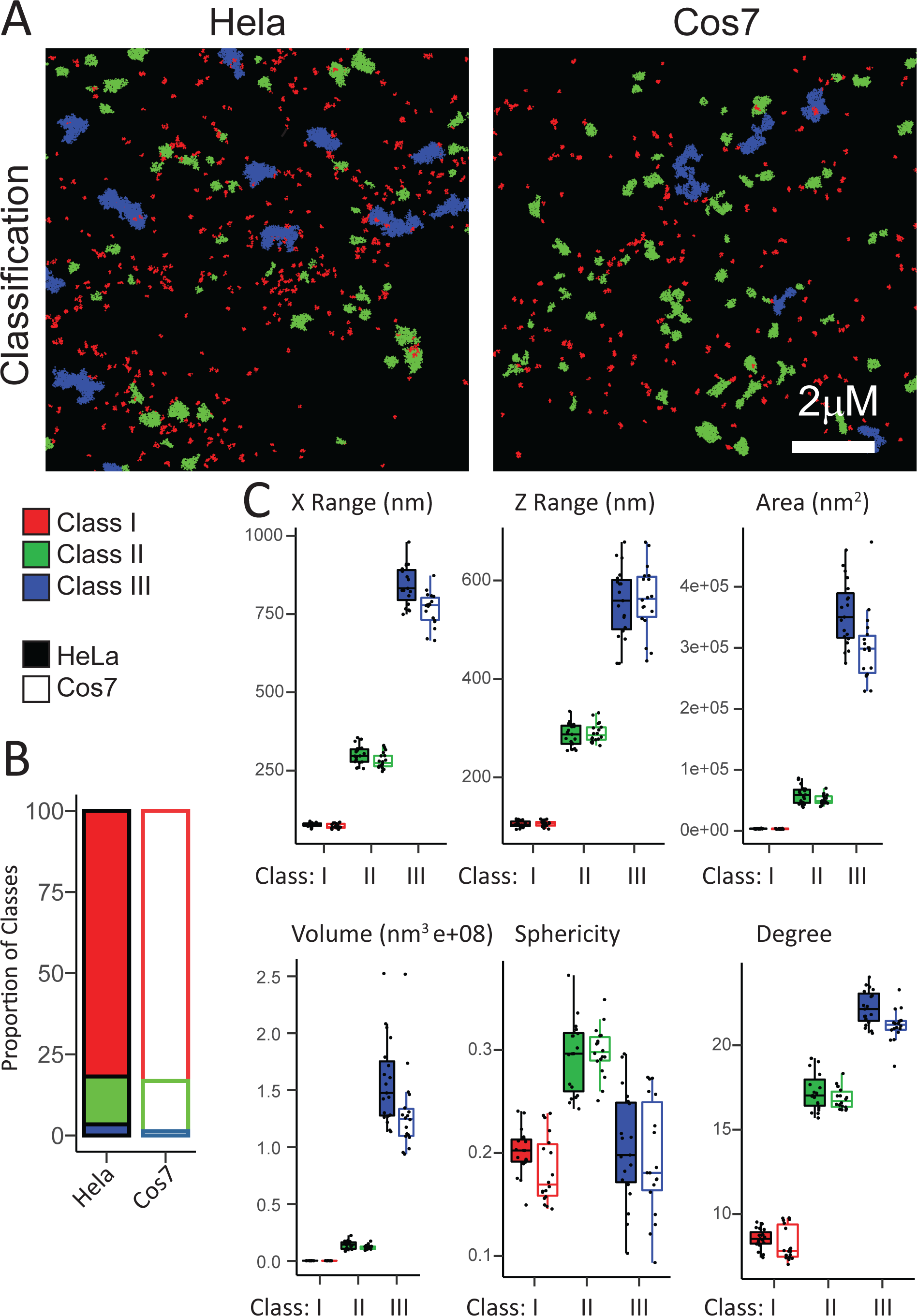
SuperResNET quantification of clathrin labeled HeLa and Cos7 cells. (A) Representative processed images of clathrin in a HeLa and Cos7 cell. (B) Proportion of clathrin Class I (red), Class II (green) and Class III (blue) in HeLa and Cos7 cells. (C) Size, shape and network degree quantification highlight the differences in features between the three classes. Scale bar 2 µm. (n = 19 HeLa cells, n = 18 Cos7 cells from three independent experiments; two-tailed unpaired t-test; no significant difference in features between HeLa and Cos7).

2D visualization of the outer boundary as well as the 3D network for representative blobs in each class are shown in Fig. 3A. Class I blobs present small, irregular shapes consistent with clathrin oligomers [27]. 2D cross sections of Class III blobs highlight their more irregular shaped structure compared to the more spherical and less dense nodes of Class II blobs (Fig. 3A). Visualization of Class II blobs shows the presence of high degree localization in the center of the cluster. We interpret these high degree localizations to reflect low levels of background labeling not removed by filtering. The analysis was performed at proximity threshold 80 nm for clathrin pits whose inner diameter is 100-200 nm diameter as reported by EM [23, 24]. Central background blinks are therefore equidistant from peripheral clathrin labeling throughout the structure and, at proximity threshold 80 nm, able to interact with blinks on the entire inner surface of the hollow clathrin blob. Cross-sectional analysis shows that upon removal of 15.9% of the highest degree localizations (retaining average + 1STD localizations), Class II blobs are hollow (Fig. 3B).

**Figure 3.**
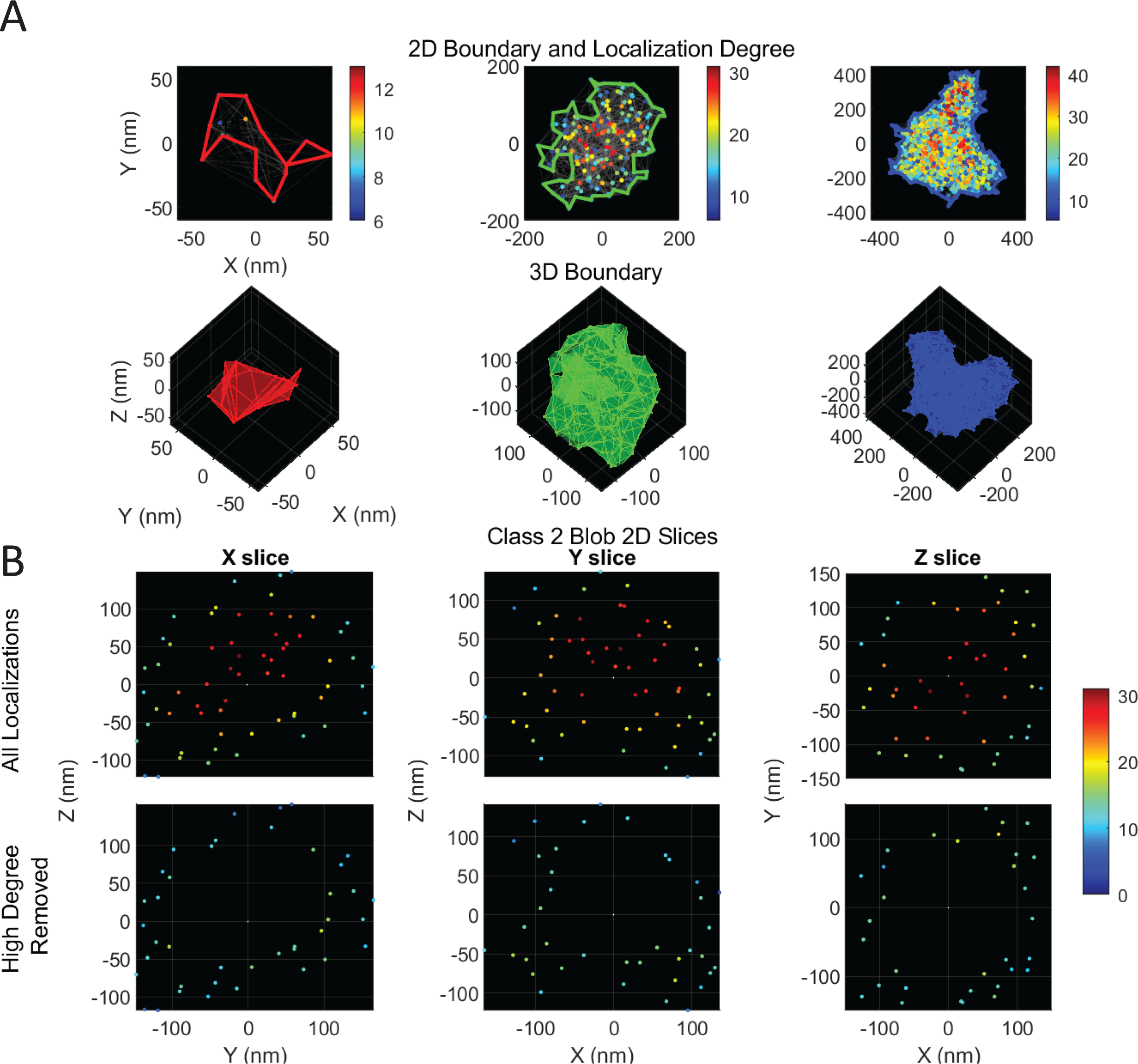
SuperResNET views of representative HeLa clathrin blobs. (A) 2D and 3D views of representative Class I, II and III blobs, extracted using SuperResNET from the HeLa cell with features closest to the average from n=19 HeLa cells. 2D views consist of nodes with colors corresponding to the node’s degree, edges generated using an 80 nm proximity threshold. A boundary box outlining the blob with a 0.5 shrink factor is used for the 2D and 3D views. (B) Cross sections of localization in the middle 10% thickness of a Class II blob are shown with nodes color-coded for degree. Bottom panels show visualization with removal of nodes with degree higher than the average plus one standard deviation.

### SuperResNET applied to MinFlux clathrin images

To generate higher resolution views of these sub-classes of clathrin pit structures, we applied SuperResNet analysis to a small dataset of SNAP-tagged clathrin light chain expressing HeLa cells labeled with Alexa647 and imaged by MinFlux with a reported axial and lateral resolution of 2 nm [12]. Mean shift segmentation classified blobs into 2 groups using K means (Fig. 4A). Classification into 2 classes effectively separates the smaller oligomers (Class 1) from the larger blobs (Class 2) including hollow pits (Fig. 4A).

**Figure 4.**
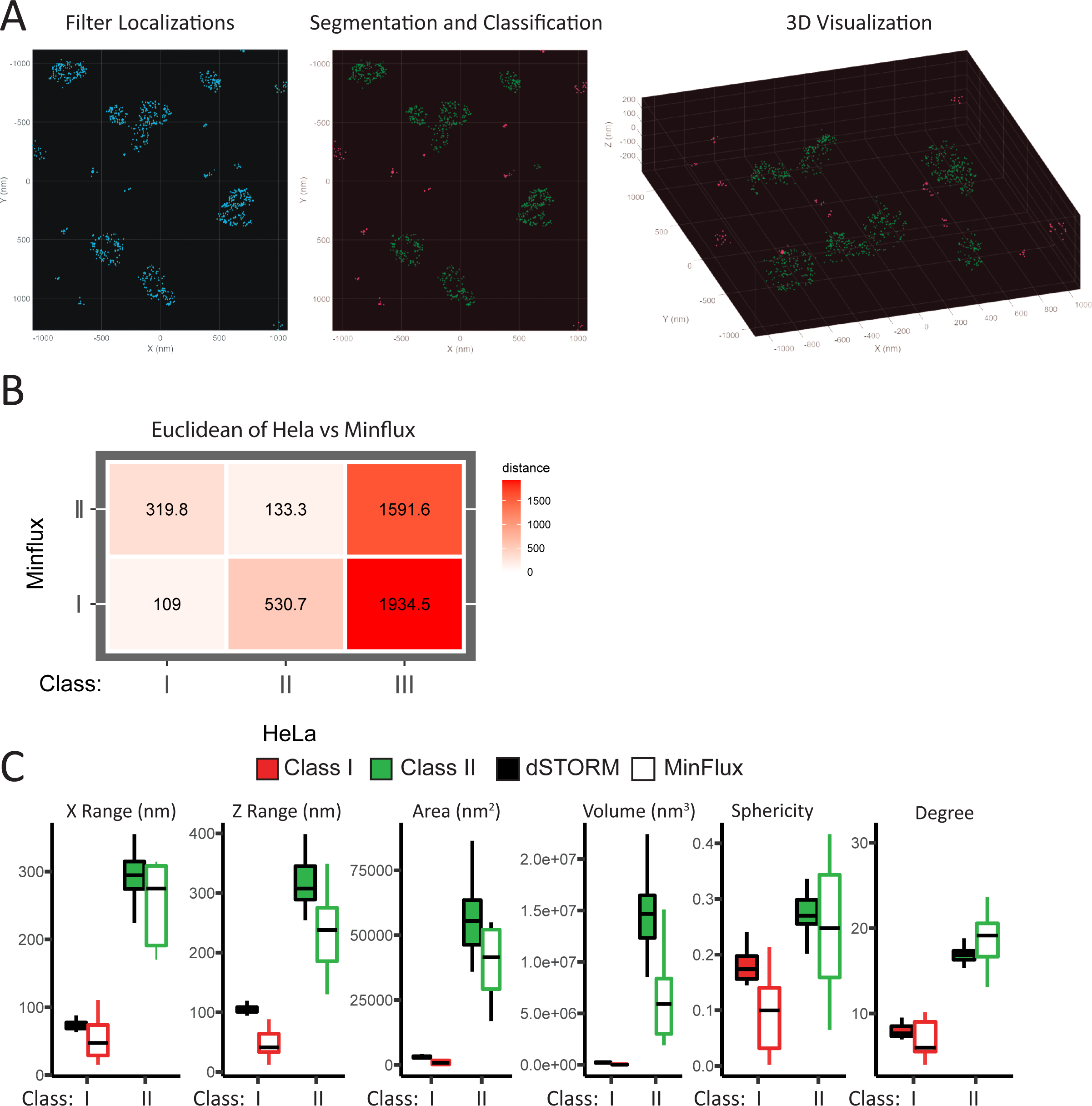
SuperResNET processing of MinFlux clathrin light chain labeled HeLa cells. (A) Point cloud representation of SNAP-tag clathrin light chain in HeLa cells, labeled with Alexa 647 and imaged using MinFlux after SuperResNET filtering, segmentation (mean shift), and grouping (K-means) into 2 groups (Class 1: red; Class 2: green). (B) Euclidean distance of the 27 features comparing the three classes of dSTORM HeLa clathrin blobs and two classes of Minflux blobs, excluding volume and area features (C) Size and shape features comparison of Minflux Class 1 and 2 blobs compared to dSTORM HeLa Class I and II blobs (two-tailed unpaired t-test; no significant differences were observed between dSTORM MinFlux blobs of the same class).

We then compared the features of the 2 MinFlux classes with the 3 classes detected by dSTORM in HeLa cells. Euclidean analysis of 28 features (excluding volume) shows that MinFlux Class 1 features correspond best with the small oligomers of dSTORM Class I while Minflux Class 2 corresponds to dSTORM Class II pits (Fig. 4B). The limited dataset of the MinFlux ROI, does not include structures corresponding to the size and features of dSTORM Class III structures (Fig. 4B). The size features of the MinFlux blobs are significantly reduced, with a largest effect seen for Z, consistent with the improved isotropic resolution of MinFlux relative to dSTORM (Fig. 4C). Visualization of MinFlux Class 2 blobs labelled ID1-9 show a range of blobs from small flat pits (middle column) to curved (right column) and hollow (left column) clathrin coated pits (Fig. 5B, See Supp. Fig. 3&4 for rotational views of the blobs). Visualization of the individual blobs highlights the greater structural detail gained by MinFlux resolution.

**Figure 5.**
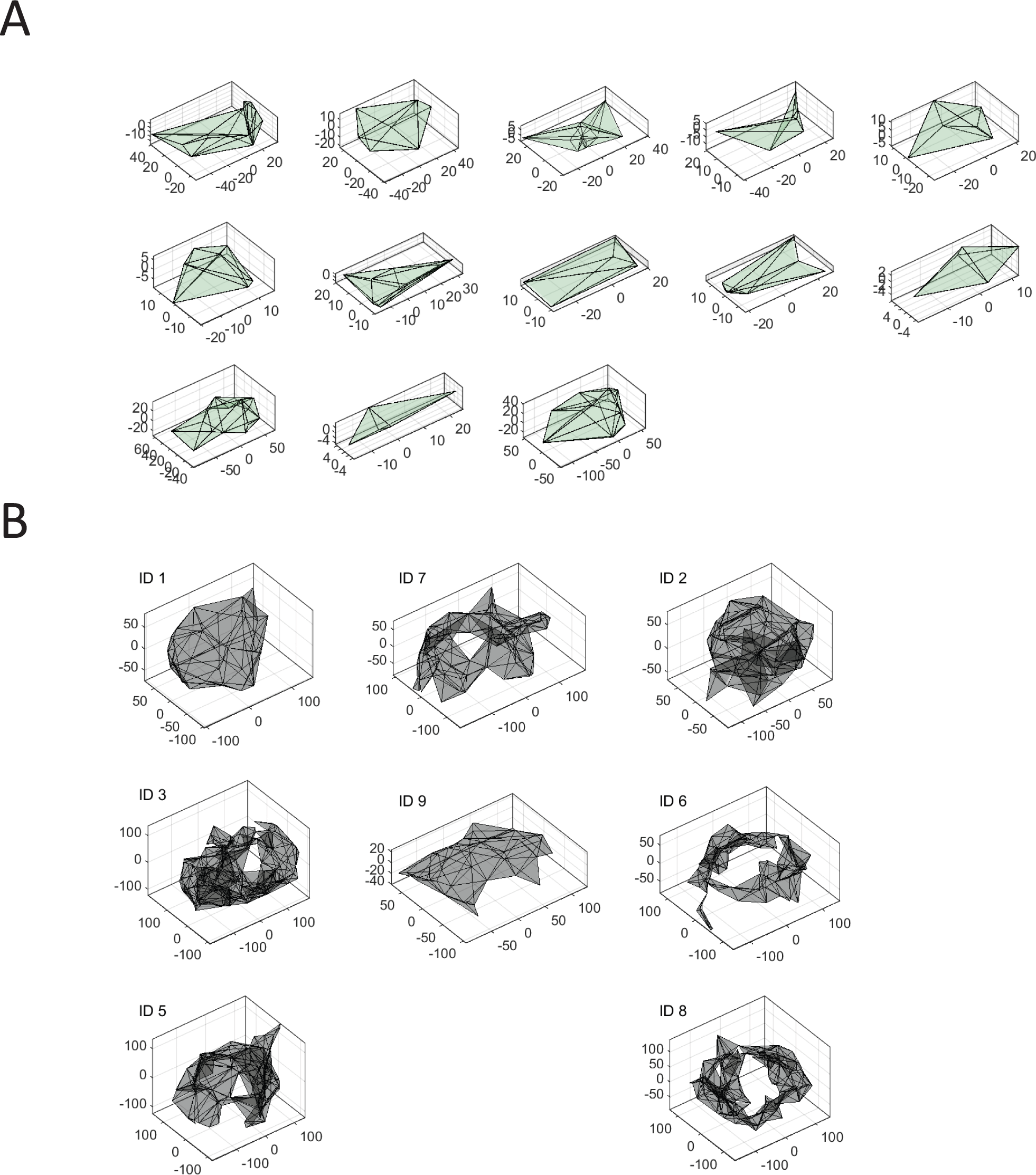
3D boundary views of Minflux groups. A boundary box with a 0.5 shrink factor was used to create a convex hull of Minflux Class 1 (A) and Class 2 (B) blobs.

### Pitstop 2 forms distinct small and dense clathrin pits

We then undertook to test whether SuperResNET could detect changes to clathrin coated pit structure induced by small molecule inhibitors of clathrin endocytosis. HeLa cells were treated with Pitstop 2, which binds to clathrin; Dynasore, which inhibits dynamin 2 scission factor; and LatA, an actin depolymerizing agent. Filtering and segmentation in DMSO, Pitstop 2, Dynasore and LatA treated HeLa cells were performed as before and blob classification was performed using k-means group centers of HeLa cells in Figure 2. SMLM images show that Pitstop 2 treated cells contained smaller pits that were not obviously hollow relative to DMSO treated control cells. Dynasore treated cells contained hollow pits similar to control while LatA treatment induced the formation of larger, hollow pits (Fig. 6A).

**Figure 6.**
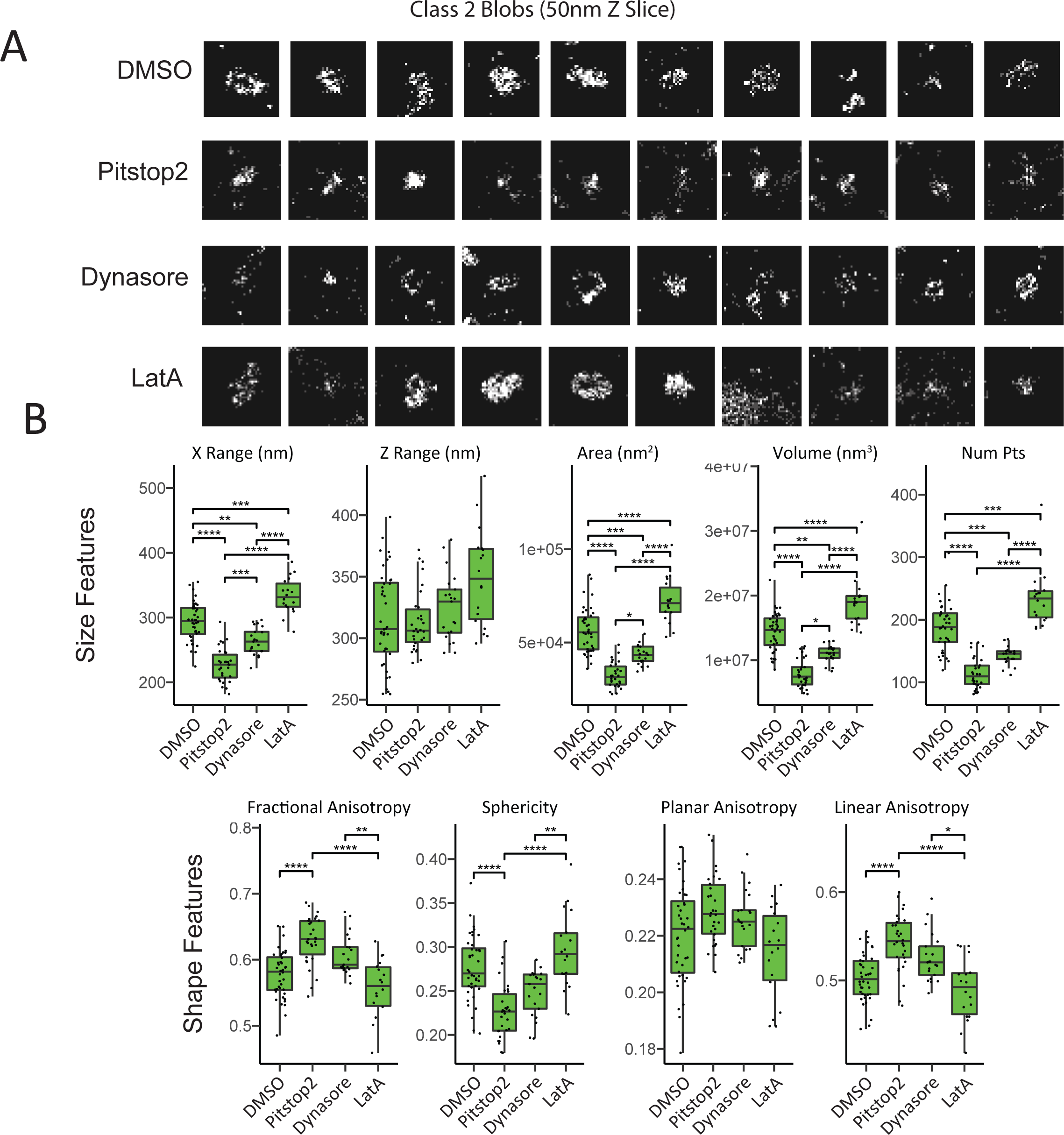
Inhibitors of clathrin endocytosis alter size and shape features of clathrin pits. (A) Representative blobs from HeLa DMSO control, Pitstop 2, Dynasore and LatA treated cells from SMLM dSTORM images. (B) Select size and shape features of Class II blobs from cells with the indicated drug treatments (n = 40 DMSO treated cells; n = 30 Pitstop 2 treated cells; n = 21 Dynasore treated cells; n = 18 LatA treated cells from six independent experiments; DMSO from six experiments, five from Pitstop 2 and three from Dynasore and LatA; ANOVA with Tukey post-test; *p < 0.05; **p < 0.01; ***p < 0.001; ****p < 0.0001).

Output of SuperResNET size features show that Pitstop 2 decreases the size of Class II blobs from 293 nm to 227 nm and Dynasore by a lesser degree to 264 nm (Fig. 6B). On the other hand, LatA increases the size of Class II blobs to 334 nm (Fig. 6B). These changes in size are more evident when reviewing blob area and volume measures and are significantly different between Pitstop 2 and Dynasore treated cells, indicating that these two drugs are differentially impacting clathrin pit structure. Variance of Class II blob size features for Pitstop 2 and Dynasore treated cells, but not LatA treated cells, was reduced relative to DMSO treated cells (Supp. Fig. 5). We interpret these results to reflect increased homogeneity amongst class II blobs in the Pitstop 2 and Dynasore treated cells, indicative of arrest at distinct stages of clathrin coated pit maturation. Dynasore is known to inhibit the final step of clathrin vesicle scission from the membrane, a step associated with condensation and reduced size of the clathrin vesicle [26]. The smaller size of Pitstop 2 treated vesicles suggests that they are arrested at an earlier stage of clathrin pit formation. Consistent with this interpretation, Pitstop 2 increased fractional and linear anisotropy and decreased spherical anisotropy of Class II blobs (Fig. 6B). In contrast, LatA treatment increased the size and variance of Class II blobs, suggesting that actin depolymerization is not arresting clathrin coated pit maturation at a specific stage nor mediating the effects of Pitstop 2 or Dynasore on clathrin pit structure. Shape features of Dynasore and LatA treated Class II blobs were not significantly different from the control, although shape features were significantly different between LatA and Dynasore treated cells (Fig. 6B).

Ternary shape feature graphs of a representative cell from each treatment group show the distribution of spherical (Cs), linear (Cl) and planar (Cp) anisotropy features for each individual blob (Fig. 7A). DMSO control, Dynasore and LatA treatment show a similar distribution of shape features, while Pitstop 2 treatment shifts the density of blobs towards lower spherical and higher linear anisotropy (Fig. 7A). Ternary graph plotting of the top 10 average dSTORM blobs based on these three shape features alongside the MinFlux blobs indicates that Pitstop 2 is shifting blobs towards the flat and more early stage MinFlux ID7 and ID9 blobs and away from the hollower, more invaginated and late stage ID1, ID3 and ID5 clathrin pits (Fig. 7B). Consistently, the 3D boundary of the top 5 representative blobs (from the 10 plotted on the ternary plot) for each treatment highlights that Class II blobs from Pitstop 2 treated cells are indeed smaller and more elongated (Fig. 7C). Rotation of the blobs shows that Pitstop 2 blobs are more irregularly shaped, while Dynasore blobs are complete spheres; DMSO control and LatA blobs are larger, spherical and bulbous (Supp. Fig. 6). These results suggest that Pitstop 2 arrests clathrin coated pit maturation at an early stage of clathrin coated pit formation.

**Figure 7.**
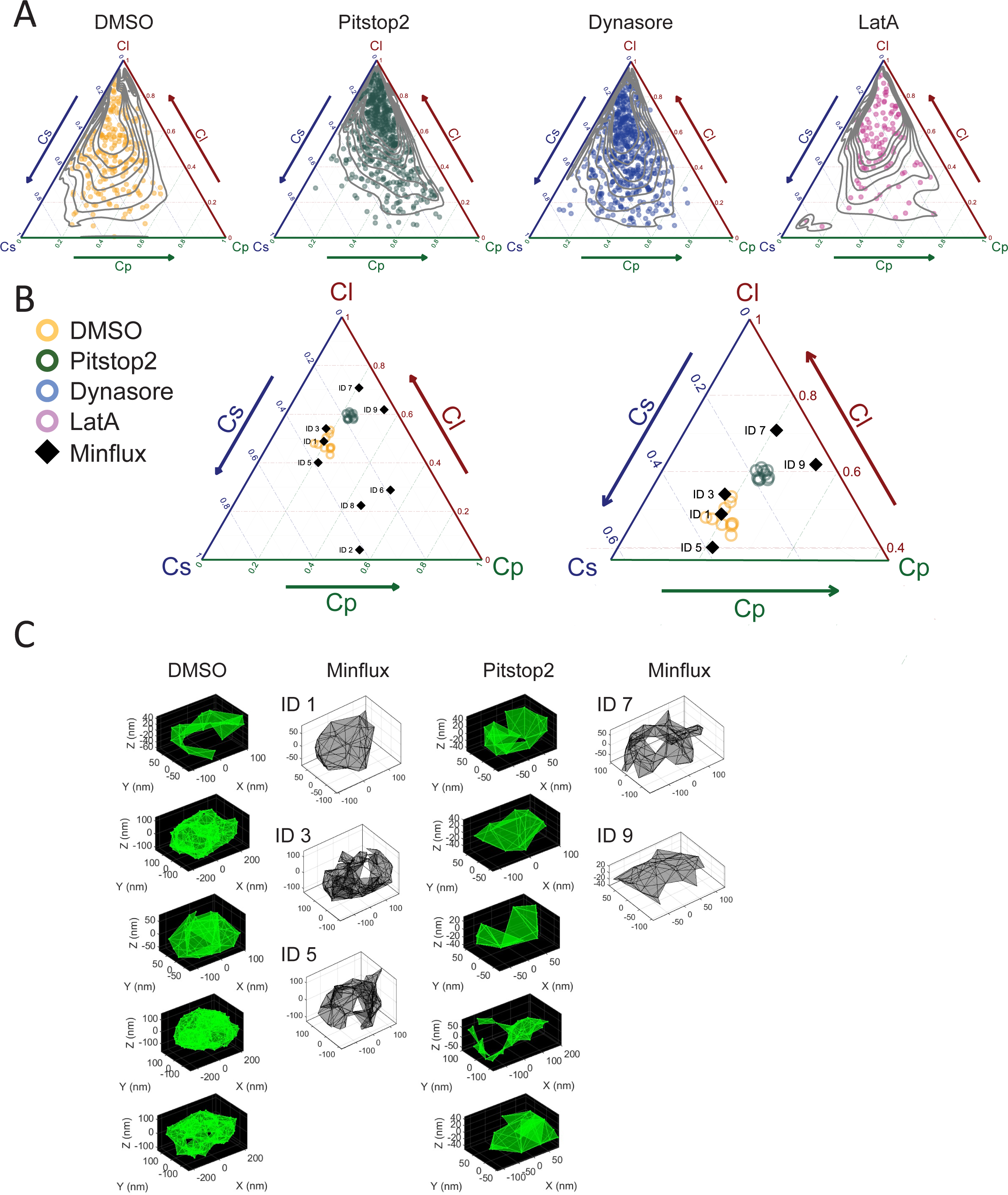
Pitstop 2 inhibits early stages of clathrin pit formation. (A) Spherical, planar and linear anisotropy (Cs, Cp, and Cl) ternary graphs of individual blobs from the top representative cells from DMSO control, Pitstop 2, Dynasore and LatA treated HeLa cells. Representative cells from each treatment were determined by selecting the cell with the closest Euclidean distance of the 27 features (with the exclusion of area and volume) for each respective treatment. (B) The top 10 representative dSTORM Class II blobs extracted from the representative DMSO and Pitstop 2 treated HeLa cells were plotted on the shape feature ternary graph with the Minflux Class 2 blobs. A zoomed in region of the ternary graph is shown to highlight the Minflux blobs that are similar to the DMSO and Pitstop 2 centres. (C) Visualization of the top 5 representative Class II blobs identified above for each treatment. Minflux blobs whose shape features align with the HeLa DMSO or Pitstop 2 blob centres on the ternary graph are grouped with the respective dSTORM blobs. 3D boundaries are drawn with shrink factor 0.5 and removal of nodes with degree higher than the average plus one standard deviation for the HeLa treatment blobs.

## Discussion

### SuperResNET: A versatile tool for the analysis of SMLM point cloud data

Here, we highlight the batch analysis properties of SuperResNET software to process SMLM data from over a hundred clathrin labeled cells, including different cell lines and drug treatments. SuperResNET is applicable to point cloud data obtained from standard dSTORM as well as higher resolution MinFlux SMLM data sets. The sensitivity of SuperResNET to structural changes in the target structures is highlighted by the detection of structural differences in clathrin pits when cells are treated with the small molecule inhibitors of clathrin endocytosis Pitstop 2 and Dynasore.

Clathrin structures include not only coated pits and vesicles but also larger plaques and smaller clathrin oligomers [28]. SuperResNET is equipped with both mean-shift and DBSCAN point cloud segmentation approaches (Fig. 1E). Mean-shift is best suited to blob-like structures and worked effectively for the MinFlux data (Fig. 4). The presence of extended structures in the dSTORM clathrin images led us to segment this data set with DBSCAN. The hollow clathrin pits present in the 3D SMLM dSTORM images were referenced to establish the DBSCAN parameters to best segment individual pits. No set of parameters could perfectly segment all the pits, with some pits adjacent to other pits or extended plaque-like regions being grouped as larger Class III structures. These large structures represented a minority of clathrin labeled structure in HeLa and Cos7 cells (Fig. 2B). The most abundant structures were small Class I oligomers, that likely correspond to plasma membrane associated clathrin triskelia and clathrin-adaptor complexes that form prior to pit formation [29]. The SuperResNET X and Y range features use the outermost localizations, giving the largest diameter of the labeled clusters. Due to the addition of a primary full-length antibody, Fab_2_ secondary and the resolution of the microscope, we expect actual structures to be smaller than reported. Class I oligomers have a diameter of ∼75 nm that is smaller than the 100-200 nm diameter of clathrin pits and vesicles [23, 24]. The Class II pits and vesicles have a blob area of 56,581 nm^2^ that corresponds closely to the previously reported 54,000 nm^2^ median surface area of clathrin pits imaged by SMLM [25]. Consistently, most hollow blobs identified by SMLM in Figure 1 are classified as Class II blobs. Class III clusters are much larger in both X and Z range (Fig. 2C) with blobs being identifiable as including multiple clusters of hollow pits or and larger clusters that could correspond to plaques (Supp. Fig. 1).

To further characterize the blob classes, we utilized a MinFlux microscopy clathrin dataset in which SNAP-tagged clathrin light chain was imaged in HeLa cells with 2 nm isotropic resolution [12]. Applying the same SuperResNET feature analysis used for the HeLa dSTORM data generated two groups; Euclidean similarity analysis showed that these two groups were most similar to Class I and Class II clathrin blob of HeLa cells. As expected, the MinFlux Class 2 blobs had smaller size features than dSTORM Class II with similar shape features. SuperResNET representation of larger Class II-matching blobs in the small MinFlux dataset of clathrin pits shows the presence of flat and hollow pits as well larger vesicles (Fig. 5, Supp. Fig. 4). Classification and quantification of the MinFlux dataset demonstrated comparable features with the clathrin pits in our dSTORM dataset, highlighting the robustness of SuperResNET feature analysis across imaging platforms using either DBSCAN or mean shift for clustering of blobs.

Other approaches for the quantitative analysis of clathrin pits in SMLM have been applied with different trade-offs. One approach has been to perform manual segmentation with 2D area and sphericity measurements using traditional thresholding methods on fluorescence images by converting the SMLM data to a pixel representation for analysis [30]. This is a simple method to apply and use available tools for fluorescence-based image analysis, however pixel representation reports indirectly on SMLM data. Another approach, the application of a maximum-likelihood model fitting of clathrin pits [25], provides an elegant model for the maturation of clathrin pits to vesicles designing different models of clathrin endocytosis and fitting the SMLM data onto the models. While model fitting has the benefit of greater interpretation for clathrin endocytosis, SuperResNET utilizes a model free approach that can be applied to various SMLM data to identify and quantify structures without prior knowledge of the structures we expect to find. Indeed, previous application of SuperResNET to caveolin-1 detected novel scaffold structures in addition to caveolae and demonstrated that expression of a mutant caveolin-1 in the caveolin scaffolding domain altered structure of the detected caveolae and scaffolds [15–17].

### SuperResNET detects drug-induced changes to clathrin molecular structure in situ

Here, we demonstrate that SuperResNET can also detect changes to molecular structures in situ due to targeted drug treatment. SuperResNET detects and differentiates molecular changes in clathrin pits in response to inhibition of clathrin endocytosis with three different inhibitors: Pitstop 2, whose specificity as a clathrin inhibitor has been challenged [31]; Dynasore, that inhibits dynamin mediated scission of clathrin coated vesicles from the plasma membrane, and whose specificity has also been challenged due to its destabilization of F-actin [22, 32]; and LatA, an actin depolymerizing agent that inhibits clathrin endocytosis [33]. Pitstop 2 and Dynasore treatment reduced the area and volume of Class II pits and vesicles and the variance of the blobs, with Pitstop 2 decreasing these features to a greater extent than Dynasore. The reduced variance of the Class II blobs suggests Pitstop 2 and Dynasore arrest and accumulate clathrin at distinct stages of clathrin coated pit maturation.

Decreased size of Class II clathrin blobs upon Dynasore inhibition of dynamin corresponds to the reduced diameter of clathrin vesicles prior to fission from the cell membrane, that has been detected by EM and live cell TIRF SIM [23, 34]. On the other hand, LatA increased size features consistent with prior EM results showing that actin depolymerization induces larger clathrin coated pits [33]. These results suggest that off-target Dynasore disruption of F-actin does not mediate its inhibition of clathrin endocytosis, that is more likely specifically related to inhibition of dynamin-dependent vesicle scission from the membrane. That LatA treatment did not result in a reduction in feature variance for Class II blobs suggests that actin depolymerization does not arrest clathrin endocytosis at a specific stage and rather more generally affects clathrin structure at various stages of the pit maturation process.

SuperResNET allows for flexible feature analysis. Focusing on selective shape features and extracting features of individual blobs within a cell, instead of averaging feature values at the cell level, reports selectively on shape distribution and allows for the interpretation of segmented clathrin clusters with respect to pit formation. Shape feature analysis shows that the smaller Pitstop 2 Class II blobs also present altered shape features relative to wild-type, Dynasore or LatA treated blobs. Overlaying individual MinFlux blobs on the ternary shape feature graphs showed that MinFlux blobs segregated based on SuperResNET shape features. Pitstop shape features of the ten most representative Class II blobs segregate with those of flatter, open, early MinFlux clathrin pits and away from the more round clathrin coated vesicles. Visualization of the five most representative dSTORM blobs confirms the smaller, flatter nature of Pitstop treated blobs. Pitstop clathrin structures are distinct from those induced by inhibition by Dynasore or LatA indicating that PitStop 2 inhibition of clathrin endocytosis does not occur at the membrane scission event and is not a consequence of non-specific F-actin depolymerization. As the constant curvature or constant area models of clathrin endocytosis both propose that early clathrin structures are smaller and flatter [26, 35], our data therefore argue that Pitstop 2 can inhibit clathrin endocytosis at an early stage of clathrin pit formation. Pitstop 2 has been shown by X-ray crystallography to bind to the N-terminal domain of clathrin heavy chain, which is the mechanism that was proposed for how Pitstop 2 prevented clathrin interaction with other proteins and block clathrin endocytosis [19]. However, despite expression of a clathrin mutation to the clathrin-box motif site at the N-terminus domain which could inhibits clathrin interaction with endocytic adaptors and uptake of transferrin; Pitstop 2 treatment was shown to further reduce transferrin uptake, suggesting that Pitstop 2 interaction with N-terminus is not the sole mechanism of inhibiting clathrin mediated endocytosis [31]. Here, we suggest that Pitstop2 binding to the N-terminus domain together with other off target effects of Pitstop 2 that inhibit clathrin endocytosis are reducing size and spherical shape of the clathrin protein coat to corresponding closer with earlier stages of clathrin pit formation.

By EM, Pitstop 2 treatment caused no significant change in structures of clathrin pit intermediates [19]. SuperResNET analysis of SMLM heavy chain clathrin labeled dSTORM data is therefore able to detect structural changes of clathrin pit intermediates that are not detectable by EM. This highlights the increased sensitivity that automated and precise SuperResNET measurements on large SMLM datasets can bring to structural analysis. SuperResNET analysis of SMLM data therefore provides a robust and sensitive method to detect molecular changes of molecules targeted by small molecule inhibitors in situ in the intact cell.

## Materials and Methods

### Cell Culturing andh2 Drug Treatment Reagents

HeLa and Cos7 cells were cultured in DMEM medium (Thermo-Fisher Scientific Inc.) supplemented with the addition of 10% fetal bovine serum (FBS, Thermo-Fisher Scientific Inc.) and 2 mM L-Glutamine (Thermo-Fisher Scientific Inc.). Cells were checked for mycoplasma using a PCR kit (Catalogue# G238; Applied Biomaterial). Tissue culture grade DMSO (D2650) and LatA (L5163) were purchased from Sigma-Aldrich, Dynasore (ab120192) and Pitstop 2 (ab120687) from Abcam for drug treatments. For drug treatments, HeLa cells were incubated with 20 μM Pitstop 2, 80 μM Dynasore, 1 μM LatA or DMSO control in serum free DMEM medium for 30 minutes before fixation.

### SMLM Preparation and Imaging

Cells were seeded on coverslips (No. 1.5H) that were cleaned by 1 h sonication with 1 M aqueous potassium hydroxide, 1 h sonication with ethanol and then washed with Mili-Q water. Cells incubated for 24 h before drug treatments and fixation. Cells were then fixed with 4% PFA for 15 minutes and washed with PBS-CM. Cells were permeabilized with 0.2% Triton X-100 and treated with Image-iT FX Signal Enhancer (Thermo Fisher Scientific). Blocking was performed using BlockAid Blocking Solution (Thermo Fisher Scientific). Primary antibody incubation against clathrin heavy chain (ab21679 from Abcam) was done for 12 h at 4 °C and secondary incubation with a fab_2_ antibody (Alexa Fluor 647-conjugated goat anti-rabbit; Thermo-Fisher Scientific Inc.) for 1 h at room temperature. A second fixation was performed with 4% PFA for 15 minutes and then fiducials markers were added by incubating with 0.1 µm TetraSpeck Fluorescent Microspheres (Thermo Fisher Scientific). For imaging, the coverslips were sealed on a glass depression slide with freshly prepared glucose oxidase buffer prepared by mixing 10% glucose, 0.5 mg/ml glucose oxidase (Sigma-Aldrich Inc.), 40 μg/mL catalase (Sigma-Aldrich Inc.) and 50 mM β- mercaptoethylamine (MEA; Sigma-Aldrich Inc.) into TN buffer consisting of 50 mM Tris and 10 mM NaCl. dSTORM imaging was performed on a Leica SR GSD 3D system with a 160 × oil immersion objective lens (HC PL APO 160 × /1.43) and EMCCD camera (iXon Ultra, Andor). Event lists were acquired using 15 ms exposure times and fiducial markers were used to correct for drift using Leica LasX software.

### SuperResNET Batch Analysis

Code for SuperResNET batch analysis, including documentation to process SMLM data is available at NanoScopy AI (github.com). SuperResNET GUI version can be access at https://www.medicalimageanalysis.com/software/superresnet.

A new batch analysis process for 3D network analysis has been created to allow the users to filter, segment, extract features and classify blobs across multiple SMLM datasets automatically. Iterative merge was performed at 20 nm to correct multiple blinking attributed to single fluorophores or fluorophores in proximity below the resolution limit. To filter out background, we constructed a network graph using the localizations with a proximity threshold of 80 nm and compared the node degree of the dataset to the node degree generated from a simulated network graph of randomly distributed localizations. The filtering parameter alpha was set to 2, which determines the threshold used to remove localizations in the comparison.

Segmentation was performed using DBSCAN with Epsilon 55 and Minpt 7 parameters. Classification was performed for HeLa control and Cos7 cells by using 3 groups with K-means. HeLa drug treated datasets utilized the class centers of HeLa control cells to classify the blobs from the drug treated cells. We used principal component analysis (PCA) to orient and plot the clathrin blobs in a consistent orientation where PC1 represents the highest variation dimension of the blob’s localizations, PC2 as the second highest and PC3 and the smallest. We applied Matlab’s boundary function with a shrink factor parameter of 0.5 on the representative HeLa cell blobs to connect a 3D triangle mesh encompassing all localizations as described previously [17].

