## Supplemental Figures for "Super-resolution single molecule network analysis (SuperResNET) detects changes to clathrin structure by small molecule inhibitors": Supplemental Figure Legends.docx

**Supplemental Fig 1.** Clusters of pits unable to be properly segmented by SuperResNET. Representations generated in raw SMLM image and SuperResNET segmentation and classification within clathrin labeling of a HeLa cell. Red boxes highlight clusters of closely contacting hollow pits visualized in the raw SMLM image being segmented as a single cluster and classified as a larger class III blob.

**Supplemental Fig 2.** Features between respective classes of HeLa and Cos7 blobs are similar. Euclidean distance of the 27 features (with the exclusion of area and volume) comparing the classes of HeLa clathrin blobs to Cos7 classes.

**Supplemental Fig 3.** 3D boundary representation of the PCA for Class 1 Minflux oligomers. Matlab file allows for rotation of blobs to visualize different angles.

**Supplemental Fig 4.** 3D boundary representation of the PCA for Class 2 Minflux pits. Matlab file allows for rotation of blobs to visualize hollowness and pit shapes.

**Supplemental Fig 5.** Variance of size and shape features for HeLa cells with drug treatments. Variance of selected features calculated per cell (n =  40 DMSO treated cells; n =  30 Pitstop 2 treated cells; n = 21 Dynasore treated cells; n = 18 LatA treated cells from six independent experiments; DMSO from six experiments, five from Pitstop 2 and three from Dynasore and LatA; ANOVA with Tukey post-test; *p < 0.05; **p < 0.01; ***p < 0.001; ****p < 0.0001). See Fig 6 for corresponding averages.

**Supplemental Fig 6.** Visualization of 3D boundary for representative Class II blobs. Top 5 blobs identified for each treatment from Fig 7. Matlab files allow for rotation and visualization of 3D structures.
