## Supplementary figures and images for "Super-resolution single molecule network analysis (SuperResNET) detects changes to clathrin structure by small molecule inhibitors"

### Supp1 Pits and plaques.pdf

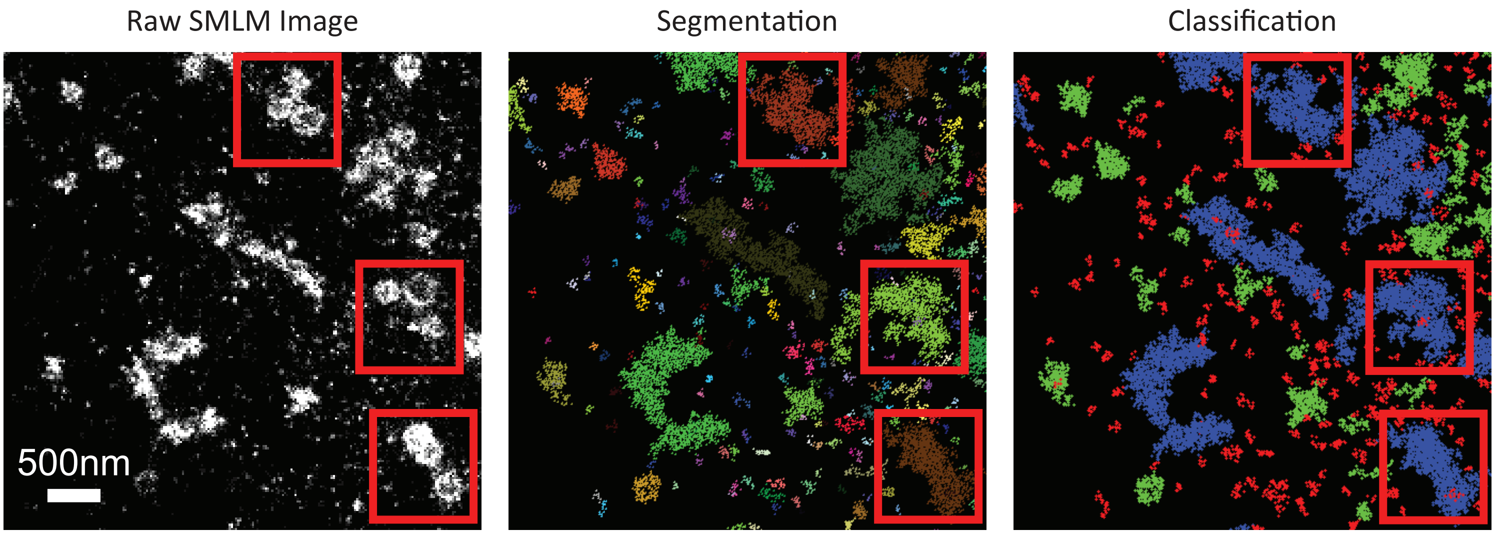


Supplemental Figure 1

### Supp2 Euclidean Distance HeLa vs Cos.pdf

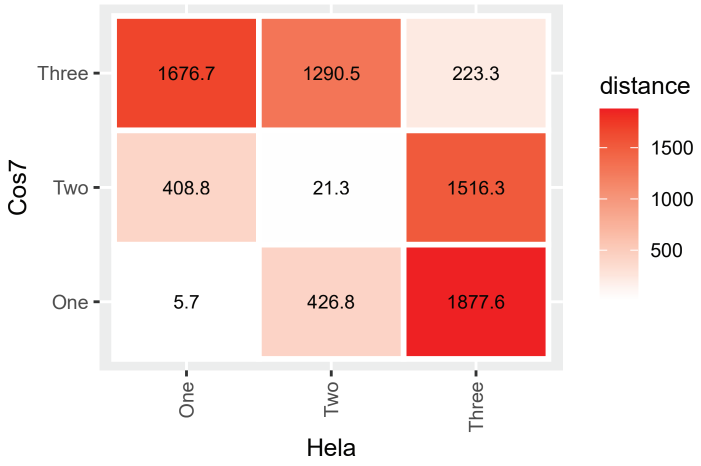


Supplemental Figure 2

### Supp5 Feature Variance.pdf

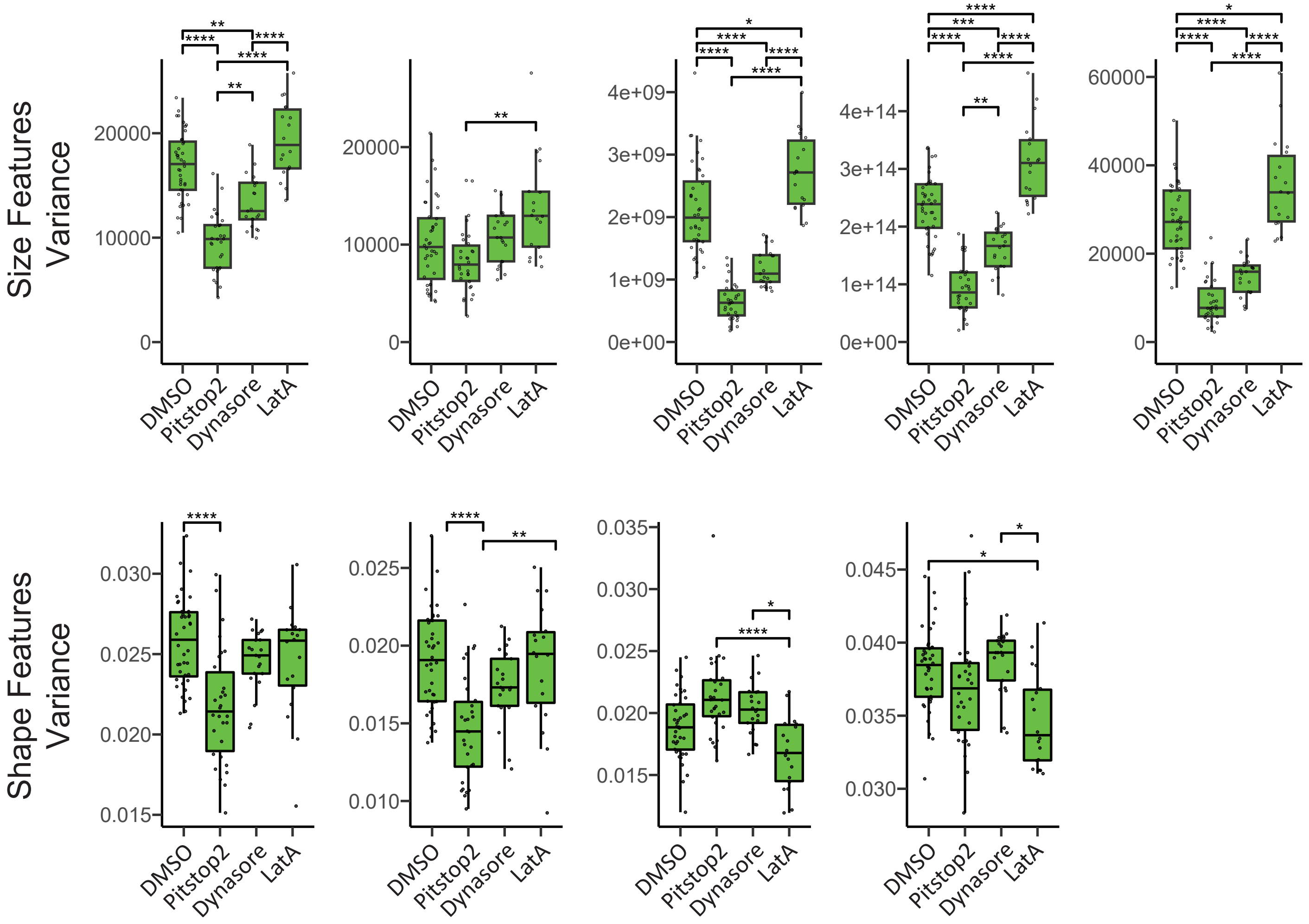


Supplemental Figure 5
